# Microbial consortia driving lignocellulose transformation in agricultural woodchip bioreactors

**DOI:** 10.1101/2024.07.11.603112

**Authors:** Valerie C. Schiml, Juline M. Walter, Live H. Hagen, Aniko Varnai, Linda L. Bergaust, Arturo Vera Ponce De Leon, Lars Elsgaard, Lars R. Bakken, Magnus Ø. Arntzen

## Abstract

Freshwater ecosystems can be largely affected by neighboring agriculture fields where potential fertilizer nitrate run-off may leach into surrounding water bodies. To counteract this eutrophic driver, farmers in certain areas are utilizing denitrifying woodchip bioreactors (WBRs) in which a consortium of microorganisms convert the nitrate into nitrogen-gases in anoxia, fueled by the degradation of lignocellulose. Polysaccharide-degrading strategies have been well-described for various aerobic and anaerobic systems, including the use of carbohydrate-active enzymes, utilization of lytic polysaccharide monooxygenases (LPMOs) and other redox enzymes, as well as the use of cellulosomes and polysaccharide utilization loci (PULs). However, for denitrifying microorganisms, the lignocellulose-degrading strategies remain largely unknown.

Here, we have applied a combination of enrichment techniques, gas measurements, multi-omics approaches, and amplicon sequencing of fungal ITS and procaryotic 16S rRNA genes to identify microbial drivers for lignocellulose transformation in woodchip bioreactors, and their active enzymes. Our findings highlight a microbial community enriched for lignocellulose-degrading denitrifiers with key players from *Giesbergeria*, *Cellulomonas*, *Azonexus,* and UBA5070 (*Fibrobacterota*). A wide substrate specificity is observed among the many expressed carbohydrate active enzymes (CAZymes) including PULs from Bacteroidetes. This suggests a broad degradation of lignocellulose subfractions, even including enzymes with auxiliary activities whose functionality is still puzzling under strict anaerobic conditions.

**Importance:** Freshwater ecosystems face significant threats from agricultural runoff, which can lead to eutrophication and subsequent degradation of water quality. One solution to mitigate this issue is using denitrifying woodchip bioreactors (WBRs), where microorganisms convert nitrate into nitrogen gases utilizing lignocellulose as a carbon source. Despite the well-documented polysaccharide-degrading strategies in various systems, the mechanisms employed by denitrifying microorganisms in WBRs remain largely unexplored.

This study fills a critical knowledge gap by revealing the degrading strategies of denitrifying microbial communities in WBRs. By integrating state-of-the-art techniques, we have identified key microbial drivers including *Giesbergeria*, *Cellulomonas*, *Azonexus*, and UBA5070 (*Fibrobacterota*) playing significant roles in lignocellulose transformation and showcases a broad substrate specificity and complex metabolic capability.

Our findings advance the understanding of microbial ecology in WBRs and by revealing the enzymatic activities, this research may inform efforts to improving water quality, protecting aquatic ecosystems, and reducing greenhouse gas emissions from WBRs.

## 1 Introduction

Elevated levels of nitrate (NO_3_^-^) and phosphorus (P) are drivers of eutrophication, habitat degradation, and loss of biodiversity worldwide ^1, 2^. Woodchip bioreactors (WBRs), also referred to as field denitrification beds (FDBs), are a technology designed to reduce non-point sources of nitrogen pollution, such as run-offs from agricultural and residential areas, by promoting microbial denitrification. The WBR is a basin filled with organic matter where woodchips typically serve as a carbon (C) source and its degradation is sustained by respiratory nitrogen oxide reduction ^3^ in two functional groups: denitrifying bacteria; these reduce NO_3_^-^ stepwise to dinitrogen (N_2_) via nitrite (NO_2_^-^), nitric oxide (NO), and nitrous oxide (N_2_O), using enzymes encoded by the genes *napAB* and *narGHI* for NO_3_^-^ reduction, *nirK* and *nirS* for nitrite (NO_2_^-^) reduction, *norBC* for NO reduction, and *nosZ* for N_2_O reduction. The other functional group is bacteria with the DNRA pathway (dissimilatory nitrate reduction to ammonium), which reduce NO_3_^-^ to ammonium (NH_4_^+^) via NO_2_^-^, and use enzymes encoded by the genes *napAB* and *narGHI* for NO_3_^-^ reduction, and *NrfAH* and *NirBD* for NO_2_^-^ reduction.

Woodchips are composed of lignocellulose, a recalcitrant co-polymer of lignin, cellulose, and hemicelluloses, and is typically degraded by the concerted action of specialized microbes, including bacteria and fungi, exploiting sophisticated enzyme systems. The most efficient degradation of lignocellulose occurs when oxygen- or hydrogen peroxide-dependent enzymes such as laccases, lignin peroxidases, and lytic polysaccharide monooxygenases (LPMOs) are utilized together with glycoside hydrolases (GHs) and other carbohydrate-active enzymes (CAZymes) ^4^. However, woodchip bioreactors are traditionally maintained under water-saturated conditions to promote denitrification (i.e., nitrate removal) which requires hypoxic/anoxic conditions. This typically leads to lower bioavailability of C from wood in WBRs ^5^, but anaerobic cellulolytic bacteria have adapted strategies for efficient cellulose degradation under such conditions. These strategies include the use of cellulosomes (large protein complexes containing multiple enzymes for efficient and simultaneous lignocellulose degradation) ^6^, and polysaccharide utilization loci (PULs) ^7^ that ensure coordinated production of “complete” enzyme sets for degradation, as well as the use of outer membrane vesicles loaded with enzymes for polysaccharide deconstructions ^8^.

Previous studies have highlighted microbial populations in WBRs involved in both C- and N-cycling (summarized by McGuire et. al. ^9^); however, only a few studies have reported on active strategies for lignocellulose degradation under denitrifying conditions within the WBRs ^9-11^. To shed more light on the expressed enzymatic repertoire of the microbial community, we have exploited an inoculum from an WBR that has been in continuous operation for three years, and used this to enrich the microbial community for further seven months on lignocellulose under denitrifying conditions in closed serum bottles, with the purpose of providing an in-depth investigation of the indigenous microbes, their capacity for C- and N-transformations, as well as their repertoires of active CAZymes for degradation of lignocellulose, while maintaining strict control of available oxygen and nitrate.

## 2 Methods

### 2.1 Samples and enrichments

The samples studied in this work originated from the surface and waterlogged subsurface of a WBR in Dundelum, Haderslev, Denmark (Figure 2A), where nitrate-rich agricultural drainage water passes through a bed of woodchips ^12^. Samples were collected in late November 2020 at two depths, i.e., 15 cm, denoted as surface-samples (S) and 60-80 cm, denoted as underwater samples (U). The WBR at Dundelum (544 m^2^) was established in 2018 with a vertical (top-down) flow design and the filter matrix consisted of 100% willow woodchips (Ny Vraa I/S, DK, chip sizes 0.4–6 cm). The wet filter matrix was 1.2 m deep and was overlain by an unsaturated woodchip layer of 30-50 cm to allow for methane (CH_4_) oxidation ^12^. In 2019-2020, total water flow to the WBR was 170 m^3^ m^-3^ yr^-1^ with a total N load of 1702 g N m^-3^ yr^-1^ and a total N removal efficiency of 46% ^12^.

Samples collected from the WBR were used directly for analysis of the WBR microbiome, as well as used as inoculum for enrichment cultures under denitrifying conditions. For these enrichments, samples from the WBR (a mix of woodchips, water and debris, ∼1 mL) was transferred to 120-mL serum bottles with 100 mg Whatman no.1 filter paper and 49 mL NRB medium (1 g/L NaCl, 0.5 g/L KCl, 0.4 g/L MgCl_2_, 0.25 g/L NH_4_Cl, 0.11 g/L CaCl_2_, 0.087 g/L K_2_SO_4_ in 10 mM phosphate buffer pH 6.0 and amended with 1 mL each of Vit-7 solution, trace element solution and Se-Wo solution, described in ^13, 14^ and 5 mM KNO_3_). The serum bottles were capped with a butyl stopper, and the headspace was He-flushed ^15^ and given a 1% O_2_ atmosphere (to provide energy for subsequently switching to a denitrifying metabolism) and incubated statically for 74 days at 17°C while monitoring the production of headspace gases as depicted in Figure 2B and described below. Thereafter, we made a subculture by taking a 1:5 dilution to new media and monitored this for another 118 days (Figure 2B, Figure S1), primarily to get rid of simple carbon substrates potentially available in the original samples and to promote growth on complex lignocellulose. The cultures were terminated at the end of this period and aliquots of 7 mL were frozen at -80°C for later meta-omics analyses.

To trace microbial growth and denitrification activity during the enrichment period, we monitored the production of gases by sampling from the serum bottle headspaces. A temperature-controlled robotized incubation system ^15^ connected to a 789A GC-system (Agilent Technologies) and a chemiluminescence NO analyzer (Model 200A, Teledyne Instruments) was used to monitor and measure the headspace concentrations of NO, N_2_O, N_2_, and O_2_. The sampled gas was replaced by an equal volume of He to maintain the pressure. Enrichment cultures were initially provided with 5 mM KNO_3_ (=250 µmol NO_3_^-^-N bottle^-1^) and N-gas production was calculated as the sum of NO, N_2_O and N_2_. Another pulse of 5 mM KNO_3_ was added when near-stoichiometric concentrations of N_2_-N were measured in the bottles.

### 2.2 Metagenomic sample preparation and data acquisition

Samples of 10 g of woodchips from the WBR were cut into pieces of 0.5 by 2 cm and shaken over night with 25 mL M9 buffer (mixture of Na_2_HPO_4_, KH_2_PO_4_, NaCl, and MgSO_4_) ^10^ at room temperature. DNA were extracted using DNeasy PowerSoil Pro Kit (Qiagen, Germany) according to the manufacturer’s protocol. For both WBR and enrichment samples, the pellet of a 4 mL sample was re-suspended before mechanical cell disruption using FastPrep24 at 6.5 m/s for 40 s followed by a 3 min incubation on ice. The homogenized lysates were centrifuged, and supernatants transferred to new tubes before subjected to DNA extraction as instructed. The quality of DNA was analyzed with Nanodrop, quantity with Qubit, and DNA degradation by electrophoresis on a 1 % agarose gel. Extracted DNA was prepared with Nextera DNA Flex Tagmentation with 350 bp and sequenced with paired-ends (2 × 150 bp) on one lane of an Illumina NovaSeq SP flow cell at the Norwegian Sequencing Center. Raw reads were trimmed, and quality control performed with TrimGalore! v0.6.6 (phred < 33, length > 20 bp) (https://www.bioinformatics.babraham.ac.uk/projects/trim_galore/), and assembled (both co- and individual assemblies) with MEGAHIT v1.2.9 (*k*-mers: 21, 29, 39, 59, 79, 99, 119, 141) ^16^. The quality-trimmed reads were mapped back to the assembled contigs using Minimap2 v2.17 ^17^ followed by Samtools v1.14 ^18^ to generate depth files for metagenomic binning. Binning was carried out with MetaBAT v2.12.1 (contig lengths > 2000 nt) ^19^ and MaxBin2 v2.2.7 (contig lengths > 2000 nt) ^20^, followed by dereplication with dRep v3.2.2 (algorithm gANi, P_ani: 0.90, S_ani: 0.97) ^21^ and taxonomical annotation with GTDB-TK v1.5.0 ^22^ and database release 202. The quality of dereplicated metagenome-assembled genomes (MAGs) were assessed with CheckM v1.2.2 ^23^ and their genomes mapped back to the high-quality trimmed reads using CoverM v0.6.1 (https://github.com/wwood/CoverM). Protein-coding genes were predicted with Prodigal v2.6.3 ^24^ and functions were predicted using InterProScan ^25^, KoFamScan ^26^ providing enzyme commission numbers (EC) and annotations from KEGG ^27^. CAZymes were predicted using the Hidden Markov Models from dbCAN ^28^ v12, downloaded from https://bcb.unl.edu/dbCAN2/ and used with the software HMMER ^29^. PULs were predicted using PULpy ^30^ which predicts the genomic location of PULs within each MAG (if present) as well as the PUL structure and components (SusC, SusD, CAZymes). Only PUL predictions containing at least one CAZyme and a SusC/D pair were considered. To quantify metabolic potential, module completion fractions (mcf) were calculated for all MAGs using the KoFamScan annotations and the R-package MetQy ^31^ to reveal the involvement of each MAG in all general biological processes (Figure S5), and in particular in denitrification and in dissimilatory nitrate reduction to ammonium (DNRA). The mcf was used to determine which metabolic pathway (denitrification/DNRA) was most likely executed by each MAG as indicated by a higher mcf-value. Read abundances of genes involved in N-metabolism were calculated using the software featureCounts ^32^ which summarizes read counts to specific genes or functional gene-groups. The phylogenetic tree of dereplicated MAGs was created with Phylophlan v3.0.60 (-- min_num_markers 50) based on predicted proteins.

### 2.3 16S rRNA sequencing and analysis

To analyze the diversity of archaeal and bacterial species present in the WBR, 16S rRNA amplicon sequencing of the V4 region was performed with Illumina MiSeq (2 × 300 paired-end) at DNASense ApS, Denmark. DNA was retrieved from subsurface woodchips in the WBR by re-extraction as described above. The quality of DNA was analyzed with Nanodrop and checked for DNA degradation by electrophoresis on a 1 % agarose gel. Raw reads were trimmed with TrimGalore! v0.6.6 (phred < 33, length > 20 bp). Read quality was further improved by adapter trimming (truncLen=c(240,150), trimLeft=c(20,21)). The DADA2 v1.26.0 pipeline ^33^, in R v 4.2.2, was used for denoising, merging and chimeric screening to finally produce 1406 amplicon sequence variants (ASVs). ASVs were taxonomically assigned using the Silva v138.1 by the *assignTaxonomy* function of DADA2. For diversity analyses, Phyloseq v1.42.0 ^34^ was used. Briefly, Alpha-diversity was measured in means of richness of three diversity indexes (Observed, Chao1, and Shannon). For Beta-diversity, samples were rarefied to an even depth using a sample size of 1000 with no-replacement by the Phyloseq function *rarefy_even_depth*. A Bray-Curtis distance matrix was calculated from these filtered samples and then ordinated using non-metric multidimensional scaling (NMDS). The statistical significance between the two different groups (i.e., surface, and underwater) was assessed by the permutation-based ANOVA (PerMANOVA) test ^35^ using the adonis2 function from the R package Vegan v2.6-6-1.

### 2.4 Analysis of fungal diversity

To analyze the diversity of fungal species present in the WBR, the ITS2 region (fITS2-C) amplicon was prepared, sequenced, analyzed, and taxonomically annotated at DNASense ApS, Denmark. Amplicon library preparation was successful for 11/12 samples, each achieving ≥ 8000 mapped reads. One sample (EU2) did not pass quality control and was excluded from further analysis. Each sample was scaled for 50k raw reads and sequenced using an Oxford Nanopore Technology R10.4.1 flow cell with the MinKNOW 22.12.7 software. The reads were basecalled and demultiplexed using MinKNOW Guppy g6.4.2 with the most accurate basecalling algorithm (config r10.4.1_400bps_sup.cfg) and custom barcodes. Sequenced reads were filtered for length (320-2000 bp) and quality (phred score > 15) using Filtlong v0.2.1. Quality-trimmed reads were mapped to the QIIME-formatted UNITE database (release 9.0)^36^ with 99 % identity clustering using Minimap2 v2.24-r1122 ^17^ and further processed with Samtools v1.14 ^37^. Entries containing *uncultured*, *metagenome* or *unassigned* were considered generic place holder or dead-end taxonomic entries and were replaced with blank entries. Mapping results were filtered for alignment lengths > 125 bp and mapping quality > 0.75, and low abundant OTUs (< 0.01 % of total mapped reads) were excluded.

To enable fungal detection in the later metaproteomics analysis, six genomes corresponding to the most abundant fungal species based on ITS were downloaded from the JGI Mycocosm database. As a complementary approach, eukaryotic contigs were extracted from the metagenomic co-assembly using EukRep v0.6.6, analyzed with Kraken2 v2.1.2 to retrieve fungal taxonomy, and their genomes downloaded from the JGI Mycocosm database. In total, fifteen fungal species were included based on this combined ITS- and Kraken2 analysis. Their proteomes were further annotated with InterProScan ^25^, KoFamScan ^26^, and dbCAN ^28^ as described above, and included in the database for metaproteomics. Differences in Alpha- and Beta-diversity analyses from fungal OTUs were performed following the same protocol as mentioned above for 16S sequencing.

### 2.5 Metaproteomic sample preparation, data acquisition, and analysis

For metaproteomic analyses, 850 mg of woodchips were mixed with 600 µL of lysis buffer (4% SDS), and 1 mL of enriched culture broth were mixed with 300 µL lysis buffer. The samples were kept on ice for 30 min, followed by mechanical cell disruption by beat-beating with FastPrep24 for three cycles of 60 s at 4.5 m/s. The samples were then centrifuged for 15 min at 15,000 × *g* and the supernatants were recovered, amended with 200 µL cold 80 % TCA, incubated over night at 4 °C, and centrifuged again at 15,000 × *g* for 15 minutes. The pellet was washed with 90 % acetone in 0.01 M HCl, the sample was centrifuged, and the supernatant was discarded. The pellet was air dried and re-dissolved in 50 µL of lysis buffer and DNA was sheered by water bath sonification for 10 min. The sample was further processed based on S-Trap 96-well plate digestion protocol (ProtiFi, USA) following the manufacturer’s instructions. The resulting peptides were re-dissolved in 0.1 % formic acid prior to LC-MS analysis.

Peptides were analyzed with a nano UPLC (nanoElute, Bruker) coupled to a trapped ion mobility spectrometry/quadrupole time of flight mass spectrometer (timsTOF Pro, Bruker). The peptides were separated by a PepSep Reprosil C18 reverse-phase (1.5 µm, 100Å) 25 cm × 75 μm analytical column coupled to a ZDV Sprayer (Bruker Daltonics, Bremen, Germany). The temperature of the column was kept at 50 °C. Equilibration of the column was performed before the samples were loaded (equilibration pressure 800 bar). The flow rate was 300 nL/minute and the samples were separated using a solvent gradient, a mixture of Solvent A (0.1 % formic acid in water) and Solvent B (0.1 % formic acid in acetonitrile). The gradient was from 2 - 25 % solvent B over 70 minutes, followed by an increase to 37 % over 9 minutes. For washing, the gradient was then increased to 95 % solvent B over 10 minutes and maintained at that level for an additional 10 minutes. The total run time was thus 99 minutes per sample.

The timsTOF Pro was operated in positive mode with data dependent PASEF acquisition. Compass Hystar v5.1.8.1 and timsControl v1.1.19 68 were used to control the LC-MS. The acquisition mass range was set to 100 – 1700 m/z. The TIMS settings were: 1/K0 Start 0.85 V⋅s/cm^2^ and 1/K0 End 1.4 V⋅s/cm^2^, Ramp time 100 ms, Ramp rate 9.42 Hz, and Duty cycle 100%. The Capillary Voltage was set at 1400 V, Dry Gas at 3.0 l/min, and Dry Temp at 180 ℃. The MS/MS settings allowed for 10 PASEF ramps, total cycle time 0.53 sec, charge range 0-5, Scheduling Target Intensity 20,000, Intensity Threshold 2,500, active exclusion release after 0.4 min, and CID collision energy ranging from 27-45 eV.

Protein quantification was performed with the MSFragger ^38^ v3.7 search engine within Fragpipe v19.0, using the workflow LFQ-MBR and Top-N (n=3) algorithm. The predicted protein-coding genes from the metagenomic analysis and the added fungal genomes were used as reference database (671,850 protein sequences). N-terminal acetylation and methionine oxidation were set as variable modifications while carbamidomethylation of cysteines were set as fixed modifications. One missed cleavage was allowed. Within FragPipe, Percolator was selected for PSM validation and FDR at 1% was allowed for ProteinProphet ^39^. The output was analyzed further in Perseus v2.0.10.0 ^40^. Identifications of potential contaminants and reversed sequences were removed. The same was done for protein hits containing indistinguishable proteins originating from other species, in order to remove cross-species protein-groups. The metagenomic annotations (taxonomy, MAG) as well as functional annotations from InterProScan, KoFamScan, and dbCAN databases were propagated to the metaproteomics data. Quantitative LFQ values for all proteins were summed per MAG to generate MAG protein-abundance (summed LFQ).

## 3 Results and discussion

### 3.1 Microbial diversity in the woodchip bioreactor

Microbial diversity within the WBR (Figure 2A) was investigated using 16S rRNA amplicon sequencing in triplicates at two depths (15 cm, denoted as surface-samples (S), and 60-80 cm, denoted as underwater-samples (U)) of the WBR. In total, 1,406 ASVs from 24 different phyla and 309 different genera could be identified (Table S1). The triplicates from surface and underwater samples showed at large a comparable and similar composition, indicating *Pseudomonas* as being the most abundant bacterial genera in the woodchip bioreactor, as also seen in several previous WBR studies ^41-43^, followed by *Flavobacterium*, *Aeromonas*, *Klebsiella*, *Chryseobacterium*, *Bacillus*, *Buttiauxella*, *Pseudarthrobacter*, *Amycolatopsis*, and *Bradyrhizobium* (Figure 1).

**Figure 1:**
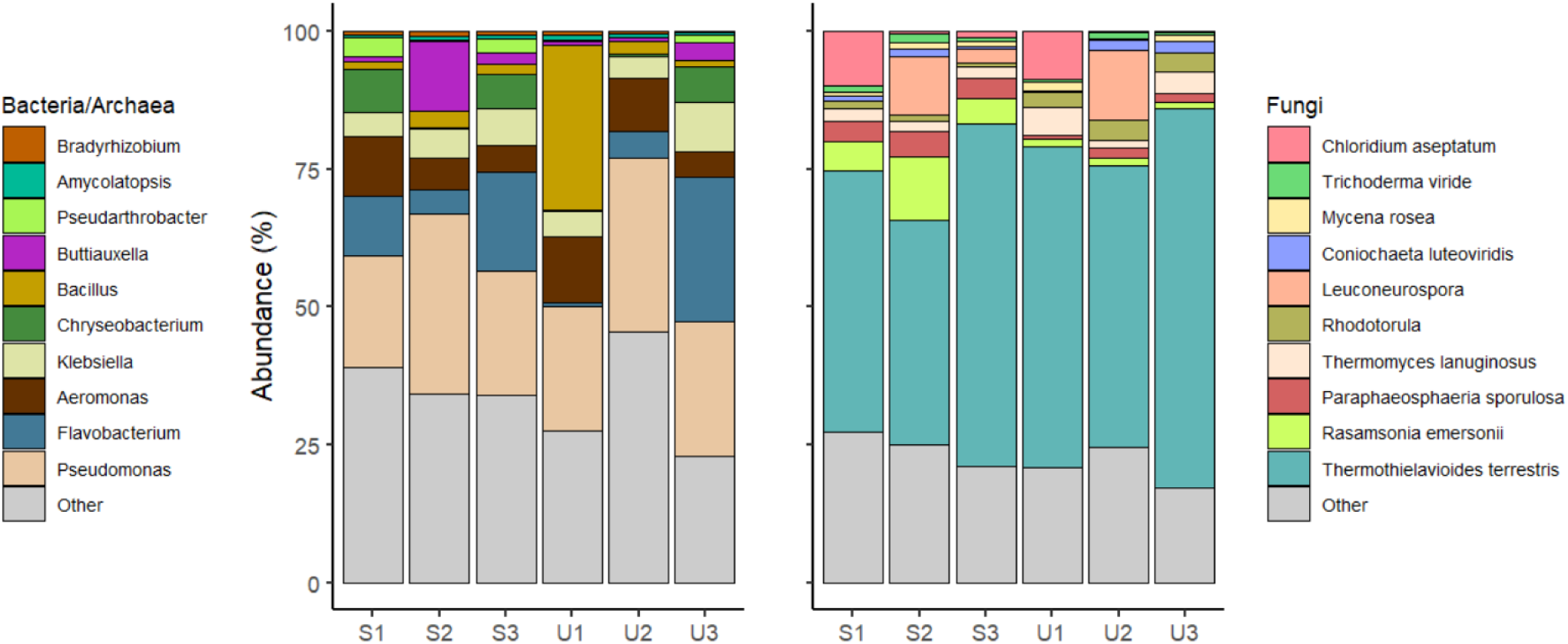
Abundance of genera within the woodchip bioreactor. The figure shows the 10 most abundant genera for bacteria/archaea (left) and fungi (right). The sample names refer to triplicates of surface (S) or underwater (U) samples in the WBR. More information about the ASVs and OTUs can be found in Table S1 and S2, respectively.

ITS sequencing was used to assess fungal diversity and identified in total 405 OTUs (Table S2) where 63% belonged to Ascomycota, 16% to Basidomycota, 11% to Chytridiomycota, and less than 3% to Mortierellomycota, Mucoromycota, Rozellomycota, and Zoopagomycota. The fungal community was by far dominated by *Thermothielavioides terrestris* (Sordariales) showing 50-60% relative abundance among the fungi, followed by *Rasamsonia emersonii* (Eurotiales), *Paraphaeosphaeria sporulosa* (<u>Pleosporales</u>), *Thermomyces lanuginosus* (<u>Eurotiales</u>), *Rhodotorula* (Sporidiobolales), *Leuconeurospora* (Leotiomycetes), *Coniochaeta luteoviridis* (Coniochaetales), *Mycena rosea* (Agaricales), Trichoderma viride (Hypocreales), and *Chloridium aseptatum* (Chaetosphaeriales). These findings are similar to prior studies on woodchip bioreactors ^9, 11^, with the exception of Heliotales which could not be detected in this study. Fungal species from the classes Chaetosphaeriales and Coniochaetales have been previously found in freshwaters ^44, 45^, while Sporidiobolales ^46^, and Corticiales ^47^ have been detected in association with plants ^46, 47^.

Due to the more permanent waterlogging in the lower area of the WBR, it is plausible that this zone has been less frequently exposed to oxygen compared to the upper parts. Consequently, the underwater zone could theoretically sustain a different set of microorganisms than those thriving on the surface. At large, only minor trends in the microbial community composition can be observed between these two locations: *Buttiauxella, Pseudarthrobacter*, and *Rasamsonia emersonii* seem to decline with depth, while *Thermothielavoides terrestris* slightly increase with depth. Analysis of Alpha-diversity showed no significant differences in measurements of Observed, Chao1 and Shannon indexes of the 16S ASVs or ITS OTUs (Figure S7). Furthermore, NMDS ordination of the Bray-Curtis dissimilarity distance showed two clusters of bacterial populations depending on the sampling site (i.e., U and S). Nonetheless, the PERMANOVA test showed no-significant difference among the clusters (Figure S7). Of note, we only had three samples in each category, making such statistical analyses less robust.

### 3.2 Relative abundances of denitrification- and DNRA pathway genes

Relative abundances of reads for denitrification and DNRA genes in the WBR ranged from <10 reads per million (rpm) to >200 rpm (Figure S2); these abundances are in the same range as previous reports of WBRs ^9^. The most abundant genes were *napA*, *narG*, *narH*, *nirB*, *norB,* and *nasA*. *nirB* is among the most abundant genes, and this seems plausible given its role in nitrite assimilation; there is ample C and nitrate in the WBR but very little NH_4_^+^ as this is not leached from soil. This may suggest that the WBR selects for microorganisms capable of assimilating nitrate, as also indicated by the relatively high value for the assimilatory nitrate reductase *nasA*. The low abundance of anammox genes is similar to what as been observed in previous WBR studies ^3, 9^.

For some of the denitrification genes (*napA* and *nirB*, and to some extent also for *napB*, *nosZ*, *nirD*, and *nrfA*), there is a clear pattern of higher abundance for underwater samples (U) compared to surface samples (S) (Figure S2). It is plausible that the more or less permanent anoxia in the deeper layer of the WBR thus favored denitrifying organisms. However, when samples from these locations were subjected to long-term enrichments under denitrifying conditions (explained in next chapter), we did not observe a higher NO_3_^-^ to N_2_ conversion rate for the underwater samples (Figure S1).

### 3.3 A microbiome enriched for lignocellulose-degrading denitrifiers

WBRs, especially those with longer operation times can be viewed as *in-situ* enrichment systems that promote microbial communities that degrade wood chips and simultaneously convert nitrate from agriculture runoff to nitrogen gases ^5, 48^. To amplify the relative biomass of microorganisms with these specific traits and allow for their in-depth characterization with meta-omics, we set up enrichment cultures on lignocellulose (wood chips and filter paper) under denitrifying conditions using the WBR samples as inoculums. These enrichment cultures showed continuous N-gas production under denitrifying conditions (Figure 2B). After 74 days, subcultures were made, and new enrichments were set up for another 118 days. At this point, the samples had converted all available NO_3_^-^ to N_2_ (Figure S1), indicating that all nitrogen reductases required for full denitrification were present within the microbial communities; likewise, the filter paper used as carbon source was fully disintegrated suggesting the presence of various lignocellulose-degrading enzymes.

**Figure 2:**
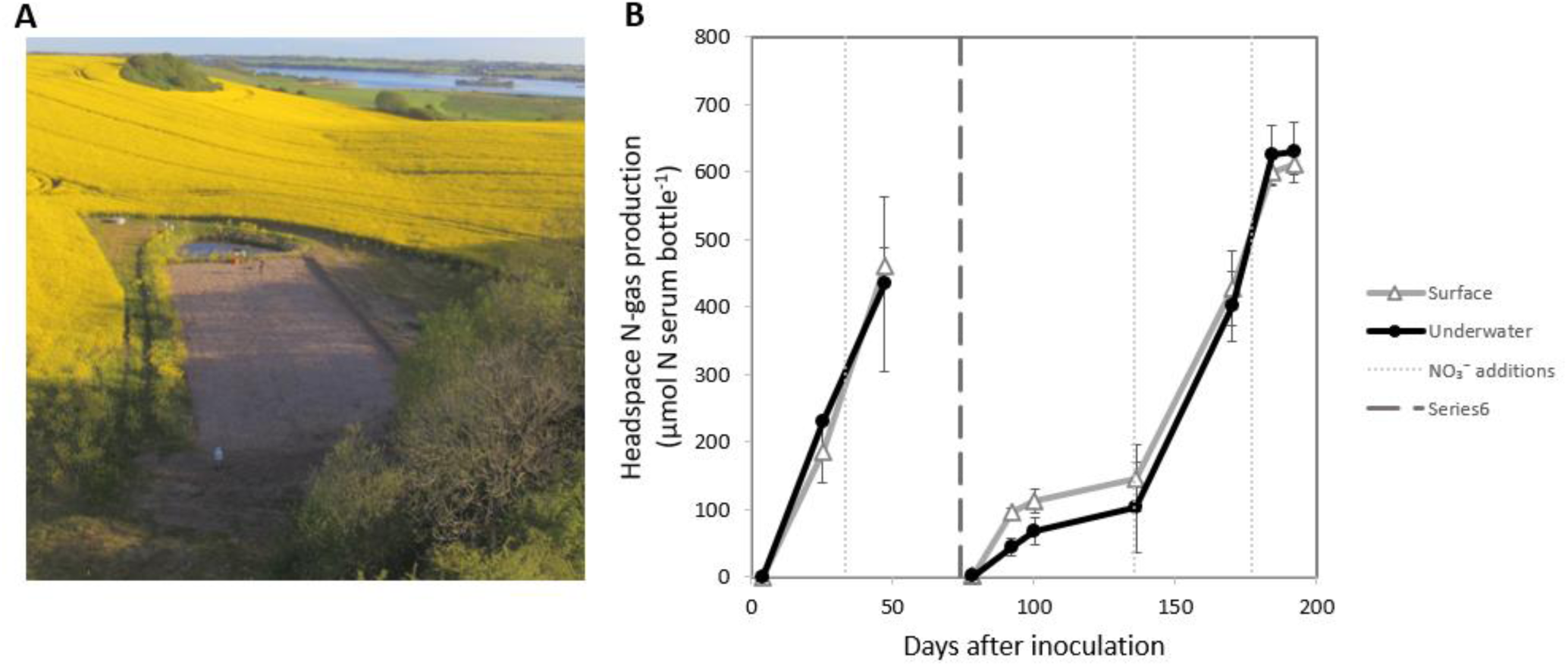
Woodchip bioreactor and enrichment cultures. **A)** Samples of woodchip filter matrix (woodchips, debris, and liquid) were collected from a field-scale woodchip bioreactor measuring 41.2 m × 13.2 m. Surface samples were collected from 15 cm depth and underwater samples were collected from 60-80 cm depth. Photo: Henning C. Thomsen, Department of Agroecology, Aarhus University, DK. **B)** Nitrogen-gas production (sum of NO, N_2_O and N_2_) in enrichment cultures with woodchips incubated in closed He-washed serum bottles under denitrifying conditions with 5 mM KNO_3_ for 74 days, followed by subculture (to dilute out simple carbon substrates from the original samples) and further enrichment for 118 days to cultivate for populations able to degrade lignocellulose under denitrifying conditions. N-gas production is averaged over three measurements; error-bars represent one standard deviation (*n* = 3). NO and N_2_O accumulated transiently and accounted for a small fraction of the N-gas produced. For details of the individual gases and incubations, see Figure S1.

Samples from both the WBR and the enriched samples were subjected for shotgun metagenomics and metaproteomics analyses. Assembly and binning of the metagenomes resulted in the recovery of 144 medium-to-high-quality MAGs, refined after binning and dereplication (Table S3). 12 MAGs were only present in the WBR in nature and did not survive enrichment cultivation, while 43 MAGs were present in both the WBR and in the enriched cultures. 89 MAGs were only detected in the enrichments. Viewing the MAG abundances, the most prominent community members in the enrichment cultures were classified as *Giesbergeria* (Proteobacteria; Bin.099 and Bin.112), followed by *Microvirgula* (Proteobacteria; Bin.111), an uncultured *Burkholderiaceae* bacterium (Proteobateria; 84.4% ANI to AVCC01; Bin.104), an uncultured *Rhodocyclaceae* bacterium (Proteobacteria; 79.2% ANI to CAJBIL01; Bin.106), *Thermomonas* (Proteobacteria; Bin125), an uncultured *Prolixibacteraceae* bacterium (Bacteroidota; 79.2% ANI to UBA6024; Bin.038), and *Cellulomonas* (Actinobacteria; Bin.009). Metaproteomics were able to detect 5,429 expressed protein groups from 141 of these MAGs (Table S4).

It was found that 92 of these 144 MAGs had the genetic potential for N-reduction, either through denitrification or DNRA (Figure 3, Figure S3). Genes required for DNRA were detected in 52 MAGs (27 complete, i.e., all genes required for DNRA were present in the genome, and 25 truncated, i.e., a fraction of the required genes were present). DNRA is catalyzed by microorganisms carrying cytochrome C_552_ nitrite reductases (*nrfA* genes) or NADH-dependent nitrite reductases (*nirB* genes), although the latter may serve as a detoxification of nitrite and NADH regeneration in fermentation of complex organic material (respiratory vs. fermentative DNRA) ^49^. For simplicity, our classification here places both pathways under DNRA. Looking at the proteomics data, no enzymes were expressed for DNRA (Figure S4), possibly due to the high nitrate-to-carbon availability benefiting denitrifying microorganisms instead ^50^.

**Figure 3:**
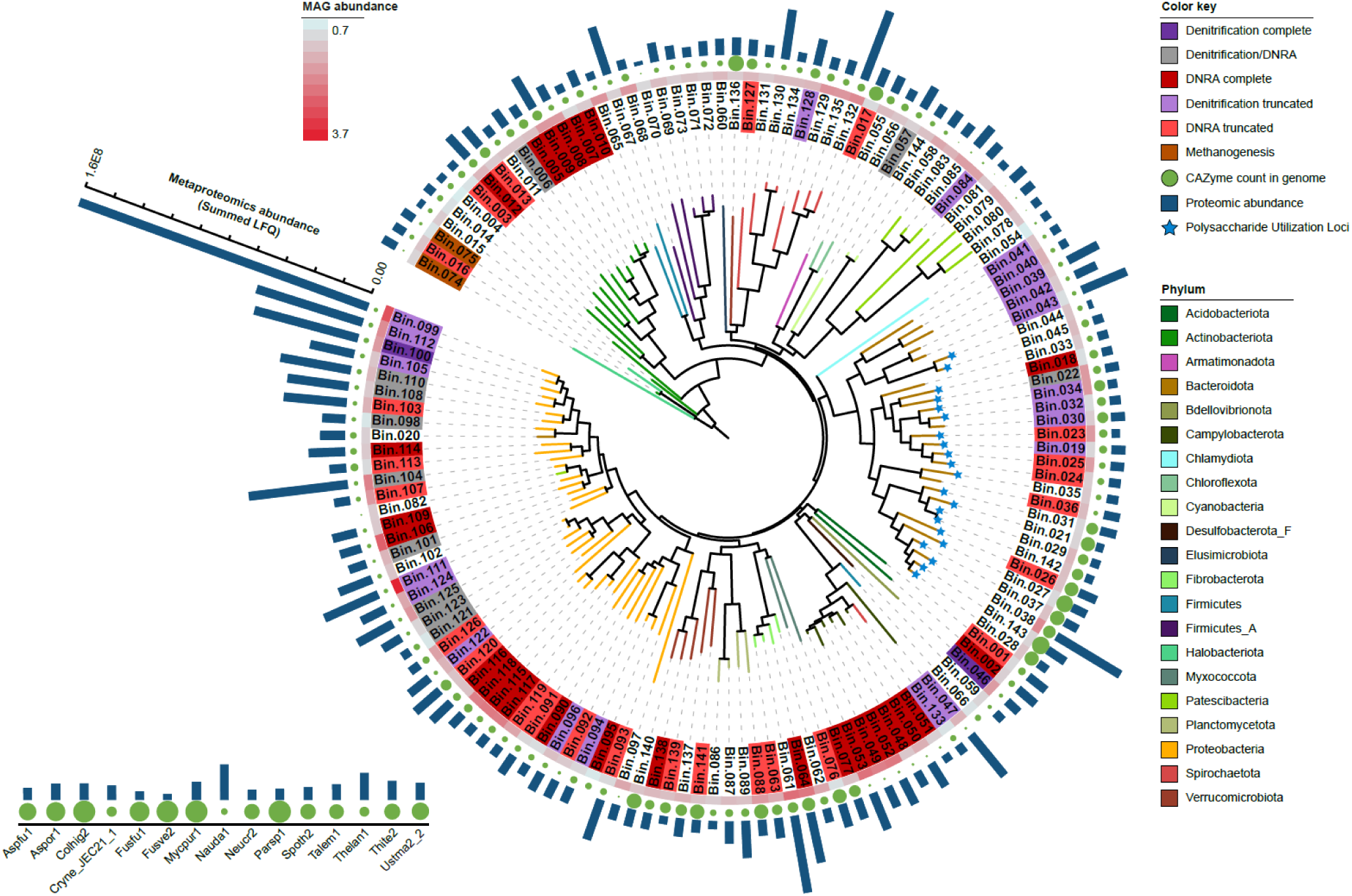
Phylogenetic placement of MAGs within the WBR and enrichment cultures. The metagenomic functional analysis of 144 reassembled MAGs from 21 different phyla show the potential for DNRA for 25 MAGs (light red) and complete DNRA for 27 MAGs (dark red). Complete full-fledged denitrification is predicted for two MAGs (dark purple) and truncated denitrification for further 27 MAGs (light purple). The classification to denitrification/DNRA were inconclusive for 11 MAGs (grey). All MAGs have genes for CAZymes in varying numbers (green circles) and polysaccharide utilization loci (PUL) could be predicted for 21 Bacteroidota (blue stars). Methanogenesis pathways can be inferred to the two archaea (brown). The circular heatmap shows metagenomic coverage, a proxy for MAG abundance, and calculated using CoverM. The bars on the outer circle represent protein abundance detected by metaproteomics. The inset to the left displays the abundance of fungi in the WBR. Abbreviations: Aspfu1 – *Aspergillus fumigatus*, Mycpur1 – *Mycena pura*, Parsp1 – *Paraconiothyrium sporulosum*, Talem1 – *Rasamsonia emersonii*, Thite2 – *Thermothielavioides terrestris*, Aspor1 – *Aspergillus oryzae*, Cryne – *Cryptococcus neoformans var neoformans*, Colhig2 – *Colletotrichum higginsianum*, Fusfu1 – *Fusarium fujikuroi*, Fusve2 – *Fusarium verticillioides*, Nauda1 – *Naumovozyma dairenensis*, Neucr2 – *Neurospora crassa*, Spoth2 – *Thermothelomyces thermophilus*, Thelan1 – *Thermomyces lanuginosus*, Ustma2 – *Ustilago maydis*.

Genes for full-fledged denitrification were detected in seven MAGs where five of them also had genes for DNRA. These included an uncultured Bacteroidales bacterium (91.2% ANI to UBA6192; Bin.022), *Bacteriovorax* (Bin.046), *Giesbergeria* (Bin.100), *Azospira* (Bin.101) an uncultured *Burkholderiaceae* bacterium (84.4% ANI to AVCC01; Bin.104), *Rhodoferax* (Bin.108), and *Acidovorax* (Bin.110). Truncated gene sets for denitrification were found in additional 27 MAGs. Expressed proteins for denitrification could be assigned to 20 of these MAGs (Figure 4, Figure S4), while none for DNRA, suggesting that denitrification is the dominant NO_3_^-^ reducing pathway in our data, as also indicated in previous WBR studies ^9, 11, 51-53^. Expressed enzymes included *napAB* (one enzyme), *narGHI* (14 enzymes), *nirK* (one enzyme), *nirS* (12 enzymes), *norBC* (six enzymes), *nosZ* (10 enzymes). The active denitrifying community, i.e., those microorganisms with expressed enzymes, is dominated by *Burkholderiaceae* as also shown previously by Jéglot et al. ^54^. Organisms with truncated denitrification pathways, lacking 1-3 of enzymes catalyzing the four steps of denitrification (NO_3_^-^-, NO_2_^-^-, NO-, or N_2_O reductases) are common in natural environments ^55, 56^. Hence, the finding of MAGs with truncated denitrification gene sets in our materials is no surprise. The propensity of a microbial consortium to emit N_2_O to the atmosphere is plausibly proportional to the relative abundance of organisms which lack N_2_O reductase. In our data, Bin.106 and Bin.111 lacks *nosZ* (Figure S4) and both show relatively high MAG abundance in the enriched cultures (Table S3); however, they have low abundance in the WBR. In the enrichment vial headspaces, all N_2_O were converted to N_2_ (Figure S1), indicating that other *nosZ*-containing denitrifiers were able to functionally compensate for the lack of this gene in these two MAGs. In an open WBR however, this N_2_O may escape as emission. Notably, under *in-situ* conditions in operating WBRs, the risk of N_2_O emissions is further controlled by factors such as the N-load, temperature and the hydraulic retention time, all which may have an effect on the capacity for complete denitrification ^53, 57^.

**Figure 4:**
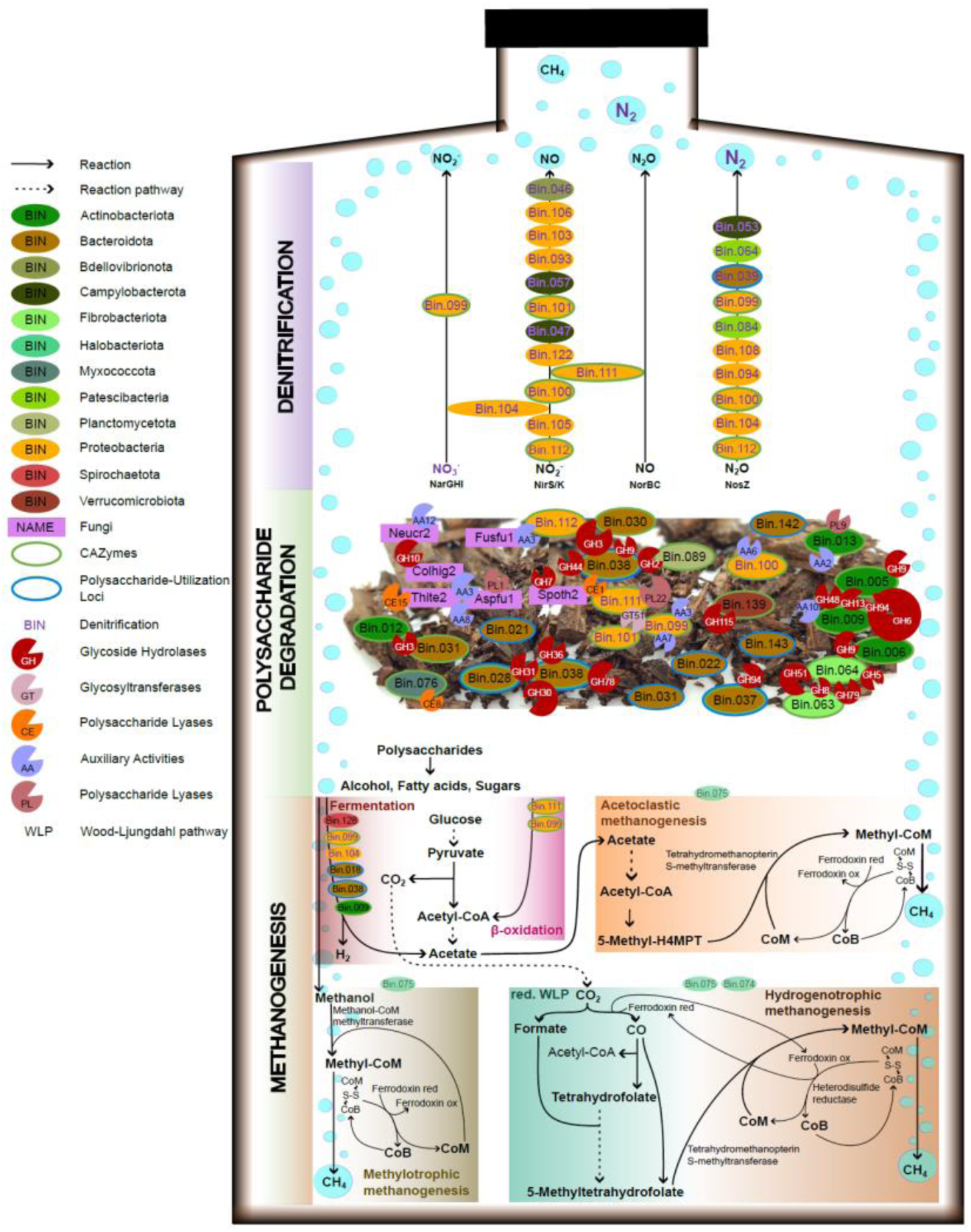
An overview of the most prominent metabolic processes in the enrichment cultures. The displayed metabolism of the microbiome is inferred using detected proteins from metaproteomics analysis and the measured headspace gases during the 7-month enrichment period. No proteins could be detected for the two archaea (Bin.074, Bin 075); however, CH_4_ production could be measured by GC suggesting that these populations were active despite being below metaproteomics detection levels. Hence, the displayed pathways for methanogenesis are inferred from metagenomics and thus indicated with a dimmed bin color. Abbreviations: Aspfu1 - *Aspergillus fumigatus*, Colhig2 - *Colletotrichum higginsianum*, Fusfu1 - *Fusarium fujikuroi*, Mycpur1 - *Mycena pura*, Neucr2 - *Neurospora crassa*, Spoth2 - *Thermothelomyces thermophilus*, Thite2- *Thermothielavioides terrestris*. More information about the N-reductases for the listed MAGs are available in Figure S3.

Diverse nitrate-reducing fungi has been reported in literature from the orders *Pleosporales*, *Heliotales*, *Hypocreales* (in particular *Fusarium* strains ^58^), *Incerae sedis*, *Thelebolales*, *Mucorales*, and *Mortierellales* ^59^, and although our metaproteomics analysis was able to detect 140 proteins from 15 different fungi (Table S4), none of these were involved in N-reduction.

Finally, the above 16S analysis showed that the genus *Pseudomonas* was dominant in WBRs (Figure 1), yet we were not able to reconstruct any *Pseudomonas* MAGs (Table S3). When investigating our list of contigs not assigned to any MAGs using Kraken2 ^60^ we observed several contigs belonging to *Pseudomonas*. Using genomes from the top-10 *Pseudomonas*-hits in our metaproteomic searches identified additional 59 expressed proteins mapping to this genus. However, none of these 59 proteins were involved in denitrification nor lignocellulose degradation; hence, despite its presence, *Pseudomonas* was not a key player in our cultures.

### 3.4 Lignocellulose is degraded anaerobically by multiple community members

While genes encoding CAZymes were found in the metagenome of all MAGs (Figure 3, green circles), the expressed enzymes were predominantly found in 24 MAGs in the enrichment samples (Figure S6). In particular, the active lignocellulose-degrading community (with CAZymes identified at protein level) included bacteria belonging to *Cellulomonas* (Bin.005, Bin.009), *Actinotalea* (Bin.006), an uncultured *Micromonosporaceae* bacterium (Bin.013), uncultured *Prolixibacteraceae* bacteria (Bin.028, Bin.037, Bin.038, and Bin.143), an uncultured *Lentimicrobiaceae* bacterium (Bin.030), *Paludibacter* (Bin.031), unclassified *Fibrobacterota* annotated as UBA5070 (Bin.061, Bin.063, and Bin.064), an uncultured *Stellaceae* bacterium (Bin.090), *Giesbergeria* (Bin.099, Bin.100, and Bin.112), *Microvirgula* (Bin.111), unclassified Polyangia annotated as Fen-1088 (Bin.076), an uncultured *Treponemataceae* bacterium classified as Spiro-10 (Bin.132), an uncultured *Opitutaceae* bacterium (Bin.139), *Amycolatopsis* (Bin.012), *Azospira* (Bin.101), and an uncultured *Phycisphaerales* bacterium classified as WQYP01 (Bin.089). In addition, CAZymes encoded in the genomes of six fungi (Table 1, Figure S6), including *Aspergillus fumigatus*, *Colletotrichum higginsianum*, *Fusarium fujikuroi*, *Neurospora crassa*, *Thermothelomyces thermophilus*, and *Thermothielavioides terrestris*, were also detected at the protein level (Table S5). Accordingly, fungal species contributing to lignocellulose degradation in WBRs have been suggested previously by Jéglot et al. ^54^

**Table 1:** Bacteroidota MAGs with PULs detected at protein level. PULs were predicted by PULPy and were included when SusC/SusD pairs were detected together with CAZymes targeting polysaccharides. 21 Bacteroidota MAGs had predicted PULs in their genomes, only PULs with components detected at protein level is shown here. More details about the predicted PULs can be found in Table S6.

| Bin | Taxonomy (Genus) | Number of PULs in genome | PUL structure with detected component in bold | Predicted target polysaccharide |
| --- | --- | --- | --- | --- |
| Bin.021 | *Paludibacter* | 19 | GH76-**susC**-susD | Fungal cell wall [α-(1→6)-mannan] |
| Bin.022 | UBA6192 (Bacteroidales) | 5 | GH13-unk-unk-unk-susD-**susC** | Starch |
| Bin.028 | UBA6024 (Prolixibacteraceae) | 16 | CE1;CBM35-GH16-unk-unk-unk-unk-CBM9-susC-susD-unk-**unk**-unk-GH28-GH92-GH92-unk-unk-GH92-GH97-unk-unk-GH78-unk-unk-susC-susD-unk-unk-GH5-GH31 | Rhamnogalacturonan I; Xylan; Galactomannan |
| Bin.031 | *Paludibacter* | 20 | susC-susD-unk-**unk**-unk-GH31 |  |
|  |  |  | GH3-susC-susD-unk-GH30-**GH30**-unk-unk-GH30-unk-unk-susC | Xylan |
| Bin.037 | UBA6024 (Prolixibacteraceae) | 18 | GH31-GH5-unk-unk-**susD-susC**-unk-unk-GH78-unk-unk-GH97-GH130-GH92-unk-unk-GH92-GH92-GH28 | Rhamnogalacturonan I; Galactomannan; |
|  |  |  | **susC**-susC-**susD**-unk-GT51 | Peptidoglycan |
| Bin.038 | UBA6024 (Prolixibacteraceae) | 14 | **susC**-unk-unk-GT2-unk-susD-**susC** |  |
|  |  |  | **susC-susD-GH36**-unk-unk-unk-GT83 |  |
|  |  |  | GH3-unk-**susC-susD** |  |
|  |  |  | **susC**-susD-GH51 | Arabinoxylan |
|  |  |  | GH42-GH43;CBM32-CE1-**susD-susC** | Arabinogalactan |
|  |  |  | **susC**-susD-GH105-unk-unk-CE1 | Rhamnogalacturonan I |
|  |  |  | GH95-unk-unk-unk-susD-**susC** | Xyloglucan |
| Bin.142 | *Paludibacter* | 19 | GH13-GH9-GH65-unk-unk-unk-**susC**-susD | Starch |
|  |  |  | **susC**-susD-unk-unk-unk-GH3 |  |
| Bin.143 | UBA6024 (Prolixibacteraceae) | 20 | GH2-GH30-unk-susD-**susC** | Xylan |

Lignocellulose, such as willow wood chips (and the cellulosic model substrate filter paper) used in this study, is a recalcitrant co-polymer of cellulose, hemicelluloses, pectin, and lignin and thus requires a multitude of enzymes with various substrate specificities for its coordinated deconstruction. The genes encoding these enzymes may be dispersed across various locations in the genome or clustered within CAZyme clusters ^61^ or polysaccharide utilization loci (PULs) ^62^.

PULs were originally identified in starch degradation (hence their name) and consist of a starch-utilization system (Sus) responsible for coordinating the activities of a set of CAZymes and carbohydrate-binding modules (CBM) involved in the degradation of a specific polysaccharide ^63, 64^. Key components of the Sus system include SusC and SusD, with SusD binding primarily to polysaccharides or cyclic oligosaccharides, while SusC facilitates the transportation of oligosaccharides into the periplasmic space of the bacterium ^65, 66^. The recruitment of specific CAZymes (including GHs, CEs and PLs) into PULs, along with the overall architecture of the PUL, appears to be connected to the glycan specificity of the PUL. More than 13,000 PULs have been identified in Bacteroidota genomes and categorized based on their specificity ^67^. In our investigation, we identified 21 Bacteroidota MAGs predicted to contain PULs. Among these, we annotated a total of 229 SusC/SusD pairs in conjunction with CAZymes, and eight of these MAGs expressed PUL components, including SusC, SusD, proteins of unknown function, or CAZymes (Table 1, Table S6). Some PULs were predicted to contain only one CAZyme, while others appeared more complex, with over 10 annotated CAZymes, often neighbored by domains of unknown function. Further examination of the annotated CAZymes in PULs having expressed components indicates that these target various plant cell wall constituents, including starch, xyloglucan, and pectin.

In accordance with the complexity of lignocellulose, using metaproteomics analysis, we detected 95 CAZymes, which could be traced back to the genomes of 24 MAGs and six fungi listed also above (Table S4, Figure S6). Most of the detected CAZymes belong to glycoside hydrolase families (GHs, 54), but auxiliary activities (AAs, 15), glycosyl transferases (GTs, nine), carbohydrate esterases (CEs, eight), and polysaccharide lyases (PLs, three), as well as proteins containing carbohydrate-binding modules (CBMs) without any identified CAZy domains (six) were also identified among the detected CAZymes (Table 2).

**Table 2:**
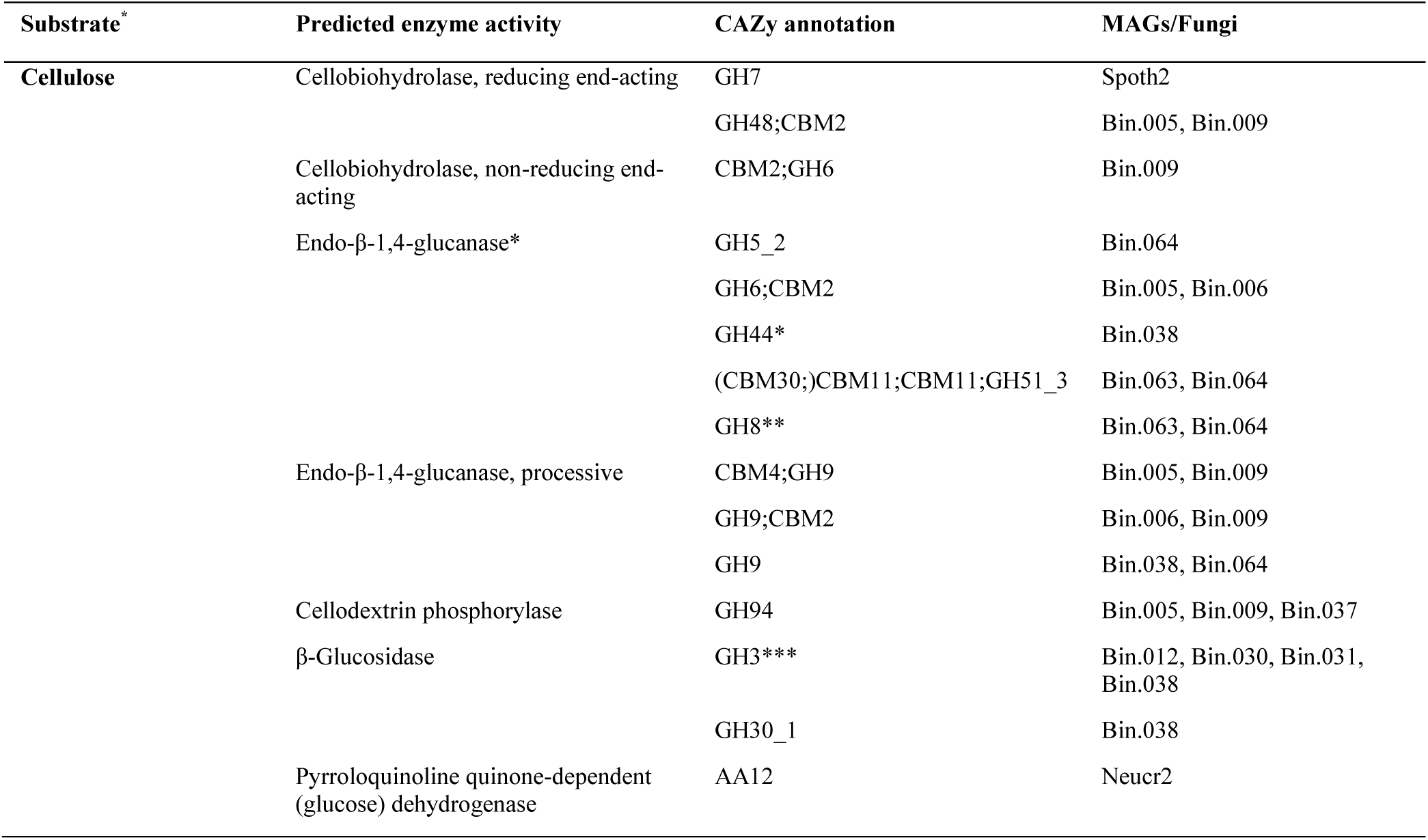
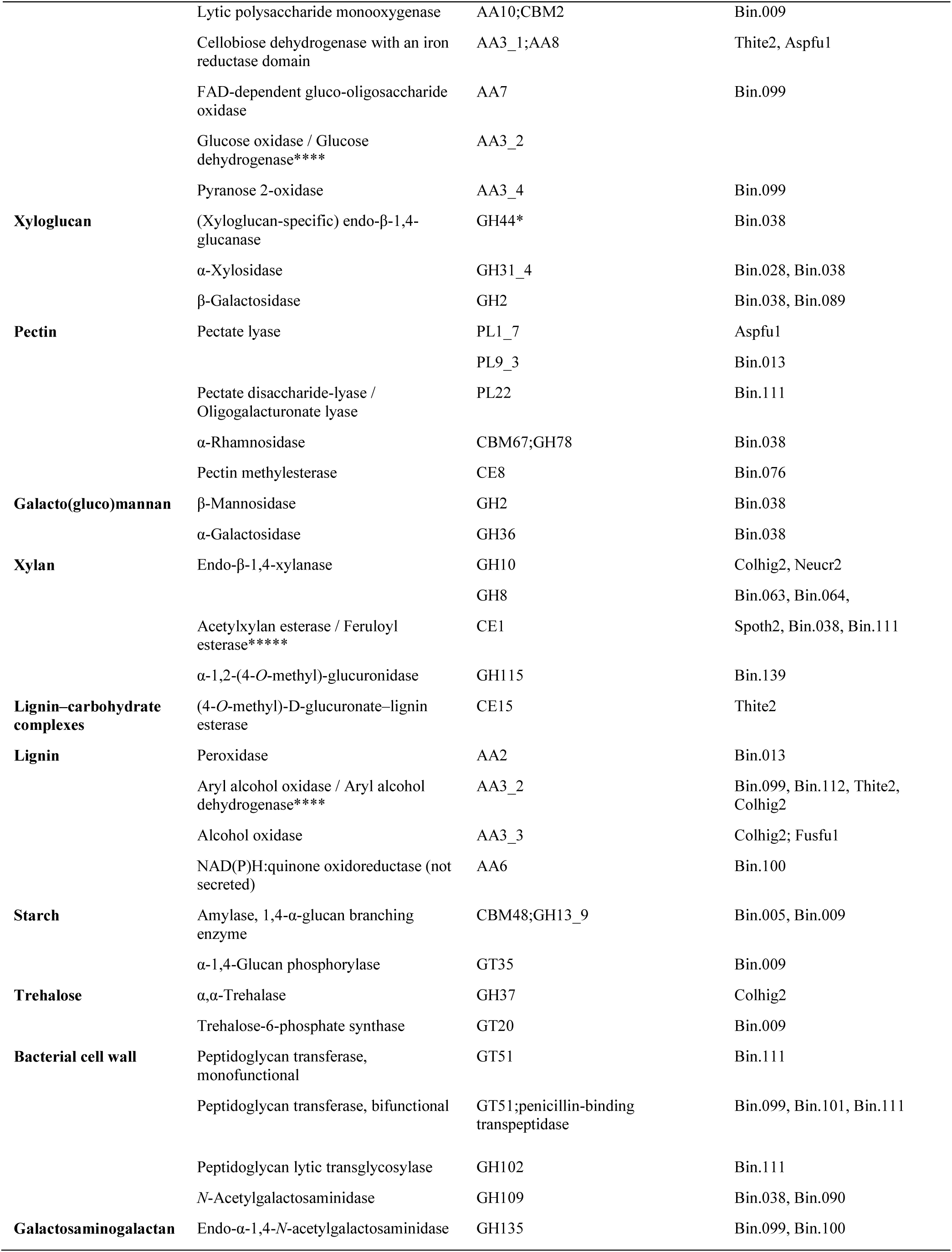
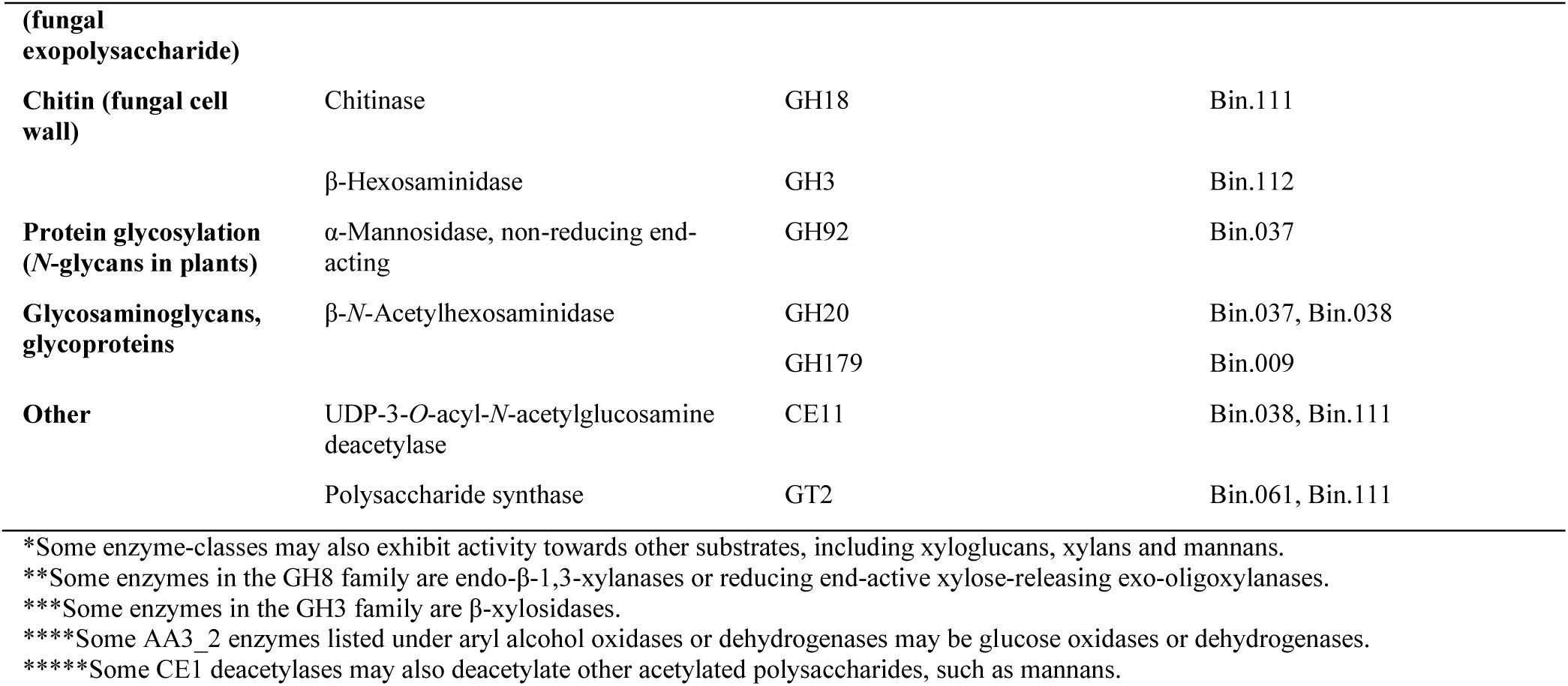
Carbohydrate-active enzymes detected at protein level that may take part in anaerobic woodchip degradation and interaction between microbial species. The table organizes the detected enzymes based on their predicted activity and target substrate and shows the CAZy modules identified in the domain structure and the MAGs/fungi expressing such proteins. Abbreviations: Aspfu1, *Aspergillus fumigatus*; Colhig2, *Colletotrichum higginsianum*; Fusfu1, *Fusarium fujikuroi*; Neucr2, *Neurospora crassa*; Spoth2, *Thermothelomyces thermophilus*; Thite2, *Thermothielavioides terrestris*; AA, auxiliary activity; CBM, carbohydrate-binding module; GH, glycoside hydrolase; GT, glycoside transferase; PL – polysaccharide lyase; CE, carbohydrate esterase.

More than half of the detected CAZymes (53) could be traced back to six bacterial species, namely: Bin.038 (fifteen) Bin.009 (twelve), Bin.111 (nine), Bin.037 (six), Bin.099 (six), Bin.005 (five CAZymes detected). Of these species, Bin.009 (a Gram-positive *Cellulomonas* bacterium) stood out as one of the main cellulose decomposers as it secreted processive glucanases. Specifically, endoglucanases (a GH9 with an N-terminal CBM4 and a GH9 with a C-terminal CBM2), which randomly cleave cellulose in the middle of the polysaccharide chain and continue releasing cello-oligosaccharides sequentially from one of the generated chain ends ^68^, and further exoglucanases (i.e., cellobiohydrolases), which target cellulose chain ends to release glucose, cellobiose or cellooligosaccharides from either the reducing end (a GH48 with a C-terminal CBM2) or the non-reducing end (a GH6 with an N-terminal CBM2 and potentially also the single-domain GH6 with a partial Fibronectin type-III module at the C-terminus) of a cellulose chain ^69^. In addition, Bin.009 also secreted an AA10 lytic polysaccharide monooxygenase (LPMO) with a C-terminal CBM2, which can oxidatively cleave crystalline cellulose in an endo fashion. Furthermore, Bin.009 seems to depolymerize cellooligosaccharides and utilize glucose as a carbon source through the phosphorolytic pathway, as it expressed a GH94 cellodextrin phosphorylase ^70^. Notably, Bin.005 and Bin.006, two other Gram-positive bacteria belonging to the same *Cellulomonadaceae* family, expressed similar enzymes for cellulose depolymerization, including a cellobiohydrolase (GH48 with a C-terminal CBM2 by Bin.005), processive endoglucanases (a GH9 with an N-terminal CBM4 by Bin.005 and a GH9 with a C-terminal CBM2 by Bin.006) and a cellodextrin phosphorylase (GH94 by Bin.005). In addition to cellulose, Bin.009 potentially degraded starch, as exemplified by a GH13_9 amylase with an N-terminal CBM48 and a GT35 α-1,4-glucan phosphorylase. Similarly, Bin.005 also expressed a GH13_9 amylase with an N-terminal CBM48.

Interestingly, Bin.038 (a Gram-negative *Prolixibacteraceae* bacterium belonging to the Bacteroidota phylum), the other major lignocellulose decomposer in the consortium, utilized cellulose-derived oligosaccharides by a different approach, secreting a β-glucosidase (belonging to both GH3 and GH30_1) to convert cello-oligosaccharides to glucose. In addition to the β-glucosidase, Bin.038 also secreted GH9 and GH44 endoglucanases (the former potentially being a processive endoglucanase and the latter being a xyloglucan-specific endoglucanase) that can depolymerize cellulose. Notably, Bin.038 secreted enzymes with activity towards a broad range of plant cell wall components, including xyloglucan, xylan, galacto(gluco)mannan, and pectin. For xyloglucan depolymerization, in addition to the family GH44 xyloglucan-specific endo-β-1,4-glucanase, cleaving β-(1→4)-linkages between the glucosyl units in the main chain, we detected a GH31_4 α-xylosidase, cleaving the xylosyl substitutions from the β-(1→4)-glucan backbone, and a GH2 β-galactosidase, cleaving galactosyl substitutions linked to the xylosyl substitutions. Furthermore, the genome of Bin.038 also contains a GH95 α-1,2-L-fucosidase, which can cleave fucosylations in xyloglucan. While the GH95 enzyme was not detected at protein level, we identified a potential PUL in the genome of Bin.038 that encodes a GH95 and the SusC component of which was identified at protein level (Tables 1, S6). Regarding galactoglucomannan depolymerization, some of the detected endoglucanases (GH44 and GH9) may potentially be able to cleave the glucomannan backbone adjacent to glucosyl units in the glucomannan backbone, while the detected GH36 α-galactosidase removes the galactosylations and the GH2 β-mannosidase and the GH3 and GH30_1 β-glucosidases monomerize the glucomannan-derived oligosaccharides. Bin.038 took part in pectin depolymerization, as exemplified by the secreted GH78 α-rhamnosidase with an N-terminal CBM67. Furthermore, Bin.038 also expressed two CE1 esterases, which most probably are xylan-specific deacetylases or feruloyl esterases.

Regarding depolymerization of xylan, we detected several secreted xylan-depolymerizing enzymes from various organisms, while we could not pinpoint any single organisms that would have played an accentuated role in xylan degradation. In particular, we detected three fungal GH10 (two from *N. crassa* and one from *C. higginsianum*) and two bacterial GH8 endo-β-1,4-xylanases (one from Bin.063 and Bin.064 each), which can cleave the xylan backbone. Moreover, we detected two CE1 esterases (one from Bin.111 and *T. thermophilus* each) in addition to the two CE1 esterases by Bin.038 discussed above. Among the xylan-active enzymes, we detected two enzymes that act on (4-*O*-methyl)-glucuronoyl groups that potentially crosslink xylan with lignin: a GH115 (4-*O*-methyl)-glucuronidase (from Bin.139), which cleaves (4-*O*-methyl)-glucuronoyl substitutions from the 2-hydroxyl groups of the xylan backbone, and a CE15 (4-*O*-methyl)-D-glucuronate–lignin esterase (from *T. terrestris*), which cleaves ester bonds formed between the carboxyl group of (4-*O*-methyl)-glucuronoyl units and phenolic hydroxyls in lignin.

It is noteworthy that despite our strict control of anaerobic conditions throughout the experiment (Figure S1), we detected a number of carbohydrate- or lignin-active oxidoreductases that require O_2_ or H_2_O_2_ as a co-substrate for their catalysis. Of the 30 species identified expressing CAZymes, one bacterial species (Bin.099, *Giesbergeria*) and four of the five fungal species (*A. fumigatus*, *C. higginsianum*, *F. fujikuroi*, *N. crassa*, and *T. terrestris*) expressed 11 of the 15 detected enzymes with known AA families and a potential role in oxidative degradation or depolymerization of lignocellulosic biomass. These included the following carbohydrate-active enzymes: a FAD-dependent AA7 gluco-oligosaccharide oxidase by Bin.099, two cellobiose dehydrogenases with an N-terminal AA3_1 dehydrogenase and a C-terminal AA8 cytochrome domain by *A. fumigatus* and *T. terrestris*, an AA12 pyrroloquinoline quinone-dependent (glucose) dehydrogenase from *N. crassa*, and an AA3_4 pyranose 2-oxidase by Bin.099. In addition to oxidation of cellulose-derived cello-oligosaccharides and glucose, many of these enzymes could potentially serve as a redox partner of bacterial AA10 (such as the AA10 LPMO by Bin.009 detected at protein level) or fungal AA9 LPMOs in cellulose depolymerization or of other AA family oxidoreductases through the generation and consumption of reactive oxygen species. Our analysis also revealed a number of oxidoreductases that are potentially active on lignin or small lignin-derived compounds, including an AA2 peroxidase (by Bin.013), five AA3_2 aryl alcohol oxidases or dehydrogenases (two AA3_2s by Bin.099 and one AA3_2 by Bin.112, *C. higginsianum*, and *T. terrestris*), and two AA3_3 alcohol oxidases (by *C. higginsianum* and *F. fujikuroi*), and an AA6 NADP(H):quinone oxidoreductase, also known as 2-hydroxy-1,4-benzoquinone reductase. Of these, at least two fungal AA3_2s (from *T. terrestris* and *C. higginsianum*) were secreted according to SignalP-5.0. For completeness, lignin can also be depolymerized by laccases (AA1) and dye-decolorizing peroxidases (DyPs) ^71^, and although many genes encoding DyPs were detected in the MAGs (e.g., four in Bin.003 and Bin.013) and fungal genomes (11 in *Mycena pura*), none of these were found expressed.

While the detection of expressed oxidoreductases remains puzzling, others have also reported the expression of similar AA enzymes in anoxia during lignocellulose deconstruction, some suggesting their potential role in Fenton chemistry ^72^. In any case, more targeted experiments are required to prove if any of these oxidoreductases would, in fact, be active during denitrifying conditions (as studied here), or if they are merely expressed due to their transcription being regulated by common inducers with e.g., glycoside hydrolases.

Last but not least, the proteomics analysis also revealed the presence of enzymes that target polysaccharides that are components of bacterial or fungal cell wall, and many of these enzymes were produced by Bin.111, a Gram-negative *Microvirgula*. The proteins active against peptidoglycan in bacterial cell wall from Bin.111 included a monofunctional GT51 and a bifunctional GH51 peptidoglycan transferase (the latter one containing a C-terminal penicillin-binding transpeptidase) and a GH102 peptidoglycan lytic transglycosylase. Furthermore, we detected a GH18 chitinase active against chitin that is present in fungal cell walls. These could be an indication of interaction between a Gram-negative bacterium and fungal species present in the WBR. Similar GH51 peptidoglycan transferases were also detected from Bin.099 and Bin.101 (one each), as well as three GH109 *N*-acetylgalactosaminidases (two by Bin.038 and one Bin.090). Corroborating the potential role of these enzymes in cross-species interactions, four of these peptidoglycan-active enzymes (a GT51 from Bin.101, a GH102 from Bin.111, and two GH109s from Bin.038) were identified to be secreted via the classical secretion pathways identified by the SignalP-5.0 server.

### 3.5 Two archaeal MAGs responsible for methanogenesis

While monitoring the gas composition in the head space of the anaerobic culture bottles, we measured an increase of methane, after all available N had been converted to N_2_ by the community (data not shown). These results are consistent with the thermodynamic preference for denitrification at low H_2_ pressure compared to methanogenesis ^73, 74^ and previous analysis of woodchip bioreactors ^51, 54^. In the WBR, we detected 26 archaeal ASVs where 21 are potentially methanogens (Table S1). All these 21 ASVs were exclusively detected in the underwater samples where more permanent anoxic conditions can be expected. *Methanosarcina* was by far the dominating genus, followed by *Methanospirillum* and *Methanosphaerula*.

Using metagenomics, we were able to reconstruct only two methanogen MAGs (Bin.074 and Bin.075). They were detected with low metagenomic coverage and few proteins (Figure 3), none of which belonged to the methanogenesis pathways; however, the MAG abundances suggest that they not only were present in the WBR but also survived in the enrichment cultures, as also evident from the gas production. However, looking at the genome reconstructions, Bin.074 is a *Methanosphaerula* population, a genus previously classified as strictly hydrogenotrophic ^75^ and herein indeed showing genes supporting hydrogenotrophic methanogenesis. Bin.075 (*Methanosarcina*) has genes for hydrogenotrophic-, acetoclastic-, and methylotrophic methanogenesis. Without detection of expressed enzymes for the methanogens, it is not possible to pinpoint which methanogenic pathways that were active under these conditions, but a likely scenario is depicted in Figure 4 and suggests that the degraded polysaccharides, now alcohols, VFA, and sugars, are being fermented into methanol, acetate and H_2_. Fermentation could be detected by expressed proteins for Bin.099, Bin.104, Bin.018, Bin.038, Bin.009, and Bin.128. Meanwhile, longer chain VFAs could be oxidized via β-oxidation (by Bin.111 and Bin.099) to acetyl-CoA, or propionyl-CoA, releasing CO_2_, which could be further converted to acetate by the acetogenic Wood-Ljungdahl pathway (Bin.009). Acetate, methanol, and CO_2_ is thus available for methanogenesis by *Methanosarcina* and *Methanosphaerula.* Due to the high level of available nitrate in the WBR and the thermodynamic preference for denitrification at low H_2_ pressure, as well as methanogenesis inhibition by N_2_O ^76^, methane emissions from the WBR to the atmosphere can generally be assumed to be low. To this end, Jéglot et al. showed only minimal methane emissions from their woodchip bioreactors ^51^.

## 4 Conclusions

This multi-omics investigation has highlighted prevailing microorganisms involved in lignocellulose transformation within WBRs. Most community members were able to respire nitrate, either through denitrification or DNRA. Still, surprisingly many MAGs did not harbor the required N-reductases and thus likely utilize an alternative respiration/fermentation strategy, potentially being strict anaerobic lignocellulose degraders, such as e.g. the *Prolixibacteraceae* bacterium Bin.038 (UBA6024). Among the MAGs capable of denitrification or DNRA, we also found several members not contributing to lignocellulose transformation but perhaps rather scavenging fermentation products from other community members, such as e.g. the *Holophaga* Bin.002. Yet, a large repertoire of CAZymes were expressed by multiple MAGs, targeting a large variety of lignocellulose subfractions, demonstrating a broad degradation of lignocellulose under denitrifying conditions. Among these denitrifying lignocellulose degraders, we find *Giesbergeria*, *Cellulomonas*, *Azonexus,* and UBA5070 (*Fibrobacterota*) to be the most abundant and active community members.

This study has presented a rich set of expressed enzymes contributing to the understanding of both nitrogen and lignocellulose transformation in WBRs and may inform efforts to optimize WBRs for improving water quality, protecting aquatic ecosystems, and reducing greenhouse gas emissions from WBRs.

### 4.1 Data availability

Raw shotgun metagenomic data has been deposited to the European Nucleotide Archive (ENA) with the accession number PRJEB73557. The mass spectrometry proteomics data have been deposited to the ProteomeXchange Consortium via the PRIDE ^77^ partner repository with the dataset identifier PXD050137. All annotated prokaryote MAGs are available publicly at https://figshare.com/projects/DENITRO-FDB/201375.

## List of abbreviations

ASV: Amplicon sequence variant
AA: Auxiliary activity
CAZyme: Carbohydrate active enzyme
CE: Carbohydrate esterase
DNRA: Dissimilatory nitrate reduction to ammonium
FDBs: Field denitrification beds
GH: Glycoside hydrolase
LPMO: Lytic polysaccharide monooxygenase
MAG: Metagenome-assembled genome
PL: Polysaccharide lyase
PULs: Polysaccharide utilization loci
WBR: Woodchip bioreactor

## Acknowledgements

This research was supported by the Novo Nordisk Foundation through grant NNF20OC0061313, by the European Research Council (ERC) through a Consolidator grant (NOD-AOL, 101125376), and by the Research Council of Norway INFRASTRUKTUR-program grant number 295910. The Galaxy server that was used for some calculations is in part funded by Collaborative Research Centre 992 Medical Epigenetics (DFG grant SFB 992/1 2012) and German Federal Ministry of Education and Research (BMBF grants 031 A538A/A538C RBC, 031L0101B/031L0101C de.NBI-epi, 031L0106 de.STAIR (de.NBI)). Parts of the computations were performed on resources provided by Sigma2 – the national infrastructure for high performance computing and data storage in Norway. We thank Arnaud Jéglot for field sampling at the WBR in Dundelum, Denmark.

